# Lis1-dynein drives corona compaction and error-correction at kinetochores

**DOI:** 10.1101/2022.03.25.485878

**Authors:** Olivera Mitevska, Pak Wing Lam, Philip Auckland

**Affiliations:** Randall Centre for Cell and Molecular Biophysics, King’s College London, London, SE1 1UL, United Kingdom

**Keywords:** Kinetochore, Mitosis, Lis1, Dynein, Spindly, CENP, centromere, microtubule

## Abstract

Mitotic cell division requires that kinetochores form microtubule attachments that can segregate chromosomes and control mitotic progression via the spindle assembly checkpoint. During prometaphase, kinetochores shed a distal domain called the fibrous corona as microtubule attachments form and mature. This shedding is mediated, in part, by the minus-end directed motor dynein, which ‘strips’ kinetochore cargoes along K-fibre microtubules towards the pole. While the main molecular players are well understood, relatively little is known about how dynein stripping is regulated and how it responds to increasing microtubule occupancy. Lis1 is a conserved dynein regulator that associates with kinetochores and is mutated in the severe neurodevelopmental disease lissencephaly. Here, we have combined loss-of-function studies, high-resolution imaging and engineered separation-of-function mutants to define how Lis1 contributes to dynein-mediated corona stripping. We show that cells depleted of Lis1 fail to fully dissemble the corona and delay in metaphase as a result of persistent checkpoint activation. Furthermore, we find that while kinetochore-tethered Lis1-dynein is required for attachment error-correction, the contribution of Lis1 to corona disassembly can be mediated by a rapidly cycling cytosolic pool. These findings support the idea that Lis1 contextualises dynein function at kinetochores to maintain corona disassembly into metaphase and prevent chromosome mis-segregation.

## Introduction

Accurate mitotic progression requires chromosomes to capture spindle microtubules and form stable bioriented attachments. To achieve this, centromeres assemble a large protein apparatus called the kinetochore. The core kinetochore is built from the constitutive centromere-associated network (CCAN) and an outer complex comprised of Knl1, Mis12 and Ndc80 (KMN) complexes, which make multiple physical contacts to centromeric DNA and microtubules (Musacchio and Desai, 2017). Metazoan kinetochores also assemble a third centromere-distal domain called the fibrous corona, named as such due to its appearance on electron micrographs (McEwen et al., 1993; McEwen et al., 1998). The corona consists of the molecular motors cytoplasmic dynein 1 (from here referred to as dynein), CENP-E and Kif2b, spindle assembly checkpoint proteins (SAC) Cyclin B, Mad1 and Mad2, dynein regulators CENP-F, Nde1, Ndel1 and Lis1, the RZZ-complex (containing Rod, ZW10 and Zwilch), the dynein adapter Spindly and CLASP proteins (Figure 1a) (Allan et al., 2020; Kops and Gassmann, 2020; Maiato et al., 2004). In the absence of microtubule attachment, the corona expands into a crescent-shaped structure that can encircle the entire kinetochore-pair (D.A.Thrower et al., 1996; Hoffman et al., 2001; Magidson et al., 2015; Pereira et al., 2018; Raisch et al., 2021; Sacristan et al., 2018; Wynne and Funabiki, 2015; Wynne and Funabiki, 2016). Formation of this supramolecular assembly is driven by a farnesylation-dependent conformational change in Spindly and Mps1-mediated phosphorylation of Rod (Pereira et al., 2018; Raisch et al., 2021; Rodriguez-Rodriguez et al., 2018; Sacristan et al., 2018). The expanded kinetochore is required for the lateral capture of microtubules, conversion of these lateral attachments into end-on attachments and the maintenance of SAC signalling (Kops and Gassmann, 2020). As end-on attachments form, the corona is disassembled into a KMN-distal remnant with distinct structural, molecular and functional characteristics. This disassembly is mediated, in part, by the minus-end directed motor dynein, which strips kinetochore cargos along K-fibre microtubules towards the poles (Auckland et al., 2020; Howell et al., 2001; Matson and Stukenberg, 2014; Wojcik et al., 2001). Recent work has shown how this stripping is negatively regulated by CENP-F bound Nde1, part of the CENP-F-Nde1-Ndel1-Lis1 (FNNL) complex (Auckland et al., 2020). The contribution of other FNNL complex members, such as Lis1, to dynein regulation at the kinetochore is less well understood.

**Figure 1:**
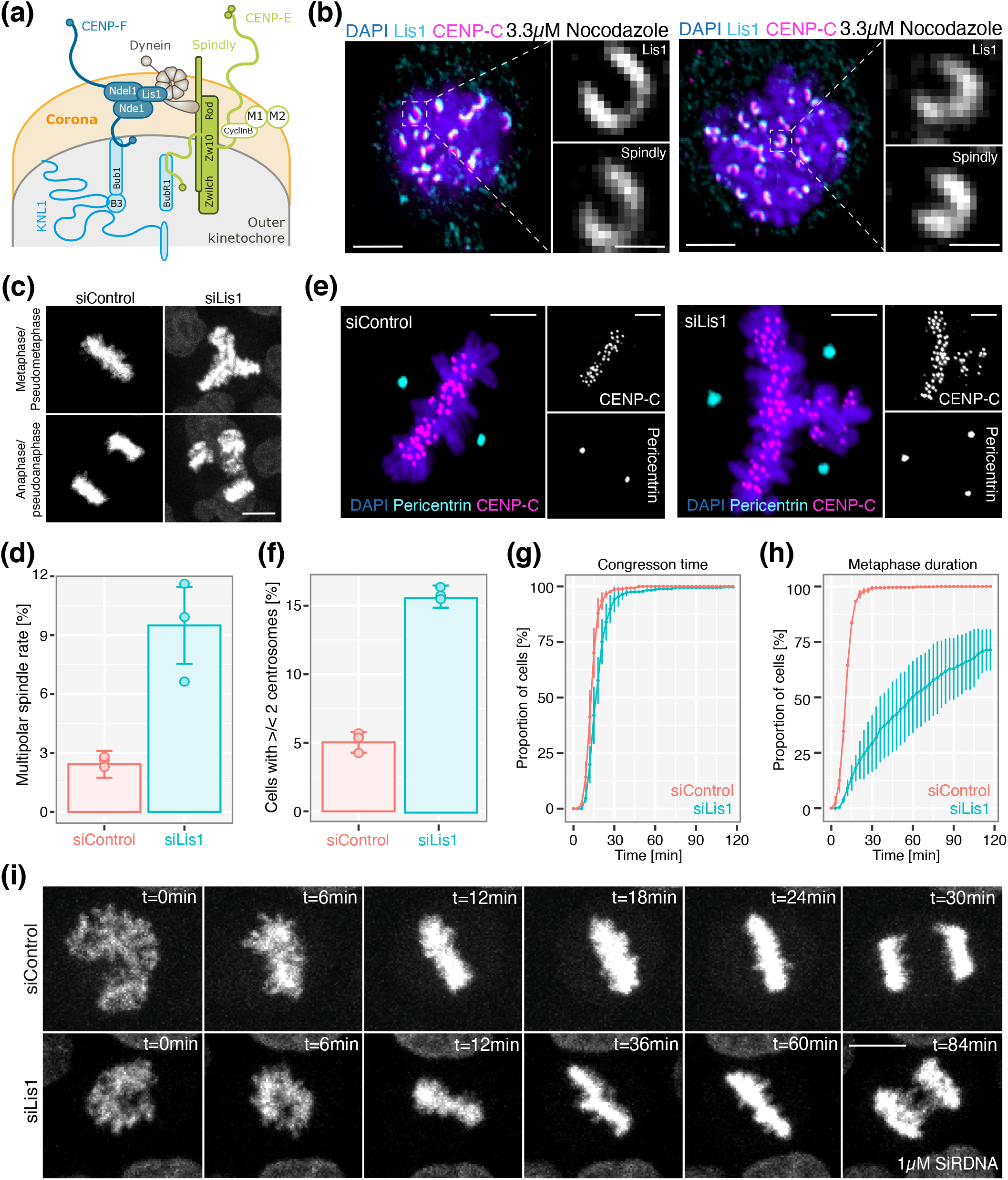
Lis1 is a corona constituent required for timely anaphase onset. (a) Cartoon schematic of major corona-associated proteins and their receptors in the outer-kinetochore. (b) Immunofluorescence microscopy images of HeLa-K cells treated with 3.3*μ*M nocodazole and stained with DAPI and antibodies against Spindly, Lis1 and CENP-C. Boxes show zooms of individual expanded kinetochores. Scale bars 5*μ*m and 1*μ*m, respectively. (c) Movie stills of bipolar or multipolar spindles in HeLa-K cells treated with either siControl or siLis1 and stained with 1*μ*M SiRDNA. Scale bar 10*μ*m. (d) Quantification of multipolar spindle rate from the live-cell imaging depicted in (c), n=3, ≥600 cells per condition. (e) Immunofluorescence microscopy images of HeLa-K cells treated with siControl or siLis1 and stained with DAPI and antibodies against CENP-C and pericentrin. Scale bars 5*μ*m. (f) Quantification of abnormal centrosome number in cells treated with siControl or siLis1, n=3, ≥550 cells per condition. (g) Quantification of congression time in HeLa-K cells treated with siControl or siLis1 and stained with 1*μ*M SiRDNA, n=3, ≥600 cells per condition. (h) Quantification of metaphase duration in HeLa-K cells treated with siControl or siLis1 and stained with 1*μ*M SiRDNA, n=3, ≥600 cells per condition. (i) Movie stills of mitotic HeLa-K cells treated with siControl or siLis1, stained with 1*μ*M SiRDNA and imaged every 3 min for 12 hr. Scale bar 10*μ*m.

Lis1 was originally identified as the gene mutated in the severe neurodevelopmental disorder Lissencephaly (Reiner et al., 1993). Lissencephaly is characterised by agyria, the absence of cerebral folding that results in brains with a smooth surface and is associated with cognitive and motor impairment, epilepsy and a shortened lifespan (Guerrini and Parrini, 2010). In lissencephalic patients, agyria is thought to be driven by defects in neural progenitor migration and division, which are required to populate the cortical plate – the precursor to the cerebral cortex (Bershteyn et al., 2017; Kato and Dobyns, 2003). Recent biochemical studies have shed light on the molecular basis of lissencephaly by establishing Lis1 a conserved activator of dynein motility (Markus et al., 2020). To summarise, Lis1 promotes the formation of activated dynein-dynactin-adapter (DDA) complexes by disrupting dynein’s autoinhibited phi state. In this ‘catalytic check value' model, Lis1 acts as a catalyst for DDA assembly while preventing reverse ‘flow’ of dynein back into the phi confirmation, which promotes both dynein localisation and cargo transport in cells (Elshenawy et al., 2020; Htet et al., 2020; Markus et al., 2020; Marzo et al., 2020; Qiu et al., 2019). In mitosis, dynein is required for spindle positioning, SAC silencing and chromosome congression. As such, perturbation of dynein or its regulators often yields complex phenotypes that confound separation-of-function analysis, a trend exemplified by current Lis1 loss-of-function studies (Monda and Cheeseman, 2018b; Moon et al., 2014; Raaijmakers et al., 2013; Tai et al., 2002). Defining how kinetochore-associated Lis1 contributes to chromosome segregation thus remains unresolved. Understanding this is key, as dysfunction of kinetochore Lis1 could contribute to the developmental defects observed in lissencephaly.

## Results

### Lis1 is a corona constituent required for SAC satisfaction

Corona proteins are defined by their inclusion into a maximally expanded domain at microtubule-unattached kinetochores (Kops and Gassmann, 2020). To test if Lis1 satisfies this criterion, HeLa cells arrested in 3.3*μ*M nocodazole (to depolymerise microtubules) were stained with antibodies against Lis1 and Spindly (Figure 1b). We found that Lis1 expanded into a crescent-shaped structure that colocalised with Spindly, thus establishing it as a component of the kinetochore corona.

While Lis1 has numerous well-defined roles, relatively little is known about its function at kinetochores. To begin addressing this, we imaged siControl or siLis1 treated cells stained with 1*μ*M SiRDNA every 3min for 12hr (Supplementary figure 1a; Figure 1c,d,g-i). Quantitative immunofluorescence revealed that siLis1 treatment reduced kinetochore Lis1 staining by 90±11% (Supplementary figure 1a). Consistent with previous reports, Lis1-depleted cells displayed an increase in multipolar divisions (from 2.5±0.5% in siControl to 9.7±2.7% in siLis1), a phenotype we confirmed in fixed cells stained with antibodies against pericentin (Figure 1e,f) (Moon et al., 2014). Nevertheless, most cells (~90%) formed a bipolar spindle and metaphase plate (Figure 1c-f). In the cells that formed a metaphase plate, the congression time was comparable to control (Figure 1g,i). In contrast, Lis1 depletion caused a penetrant metaphase delay, with only 29±12% of siLis1 cells entering anaphase within 30min compared with 99±1% of siControl cells (Figure 1h,i). To determine if this delay was SAC-dependent, we quantified the kinetochore localisation of the SAC effector Mad2 throughout early mitosis (Figure 2a,b). This revealed that kinetochores in mid-prometaphase (characterised by the presence of a forming metaphase plate with multiple unaligned chromosomes) in Lis1 depleted cells localised four-fold more Mad2 than control (Figure 2a,b). However, this increase was transient as Mad2 was undetectable at metaphase kinetochores in siControl and siLis1 treated cells (Figure 2a,b). Because undetectable levels of SAC proteins can generate a ‘wait-anaphase’ signal (Martin-Lluesma et al., 2002), we sought to determine the sensitivity of Lis1-depleted cells arrested in metaphase to pharmacological SAC inhibition (Santaguida et al., 2010). Consistent with a SAC-dependent delay, cells simultaneously exited metaphase when treated with 500nM of the Mps1 inhibitor reversine (Figure 2c). Furthermore, siLis1 treated cells incubated with reversine displayed a concentration-dependent rescue of nuclear envelope breakdown (NEB) to anaphase timing (Figure 2d,e). Here, NEB-anaphase timing decreased from 127.4±32.8min to 42.5±7.9min and 21±2.4min in siLis1 treated cells incubated with DMSO, 100nM reversine and 500nM reversine, respectively (Figure 2d,e). Taken together, these data show that Lis1 is a component of the kinetochore corona required for SAC satisfaction and timely anaphase onset.

**Figure 2:**
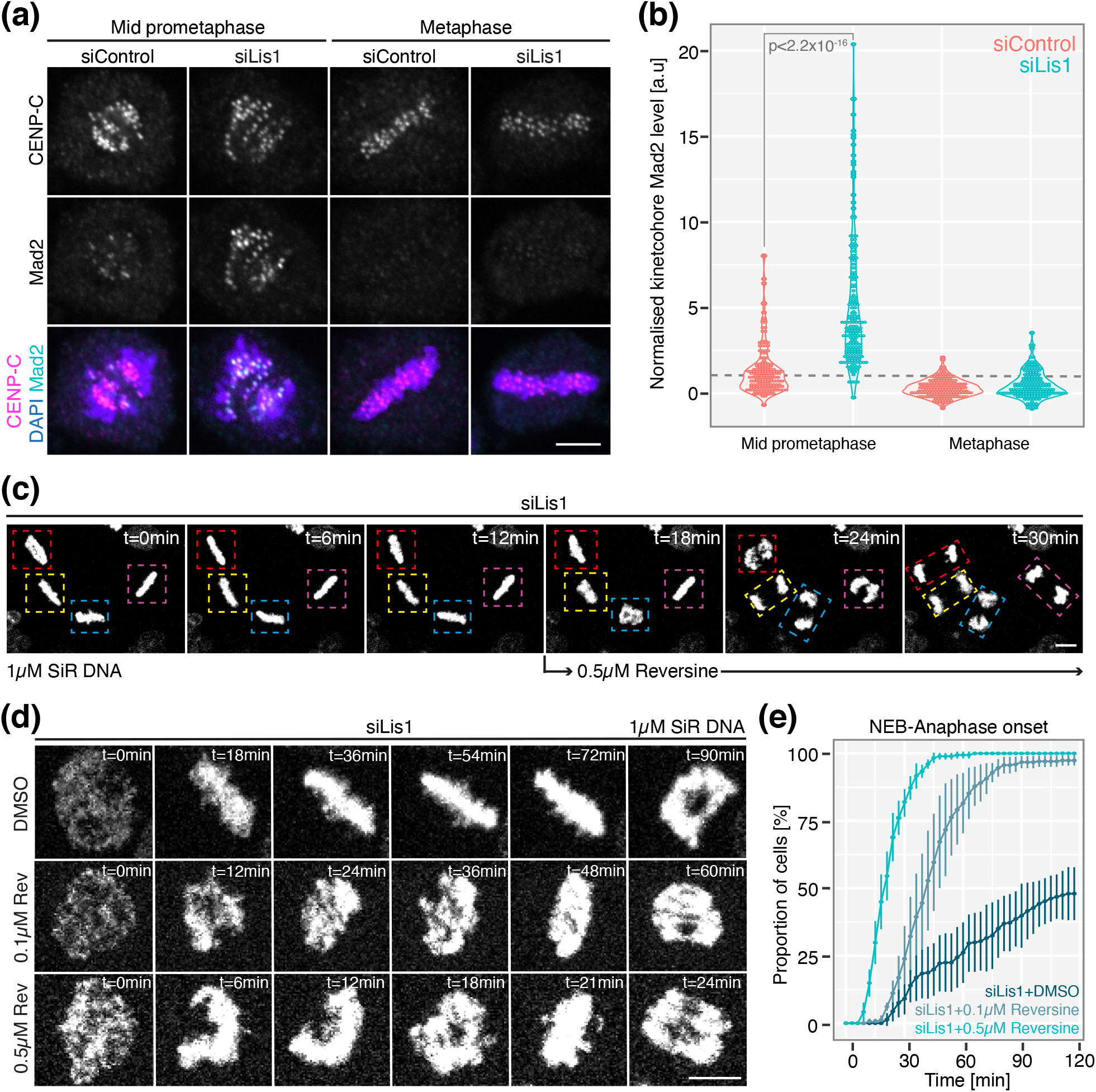
SAC-dependent metaphase arrest in Lis1-depleted cells. (a) Immunofluorescence microscopy images of prometaphase or metaphase HeLa-K cells treated with siControl or siLis1 and stained with DAPI and antibodies against Mad2 and CENP-C. Scale bar 5*μ*m. (b) Quantification of kinetochore Mad2 signal in prometaphase and metaphase HeLa-K cells treated with either siControl or siLis1, n=3, 30 cells, 300 kinetochores per condition. Dotted line represents the siControl mid-prometaphase median, to which the data were normalised. (c) Movies stills showing simultaneous exit from siLis1-induced metaphase arrest after treatment with 0.5*μ*M of the Mps1 inhibitor reversine. Cells were visualised with 1*μ*M SiRDNA. Boxes depict individual cells. Scale bar 10*μ*m. (d) Movie stills of siLis1 treated cells undergoing mitosis in the presence of DMSO, 0.1*μ*M reversine or 0.5*μ*M reversine. Cells were visualised with 1*μ*M SiRDNA. Scale bar 10*μ*M. (e) Quantification of nuclear envelope breakdown to anaphase onset time in siLis1 treated cells incubated with DMSO, 0.1*μ*M reversine or 0.5*μ*M reversine, n=3, ≥202 cells per condition.

### Lis1 is required for corona disassembly

Our previous analysis found that Lis1 depleted cells had increased Mad2 localisation to kinetochores in mid-prometaphase. To determine if this effect was Mad2-specific or reflected a broader change in corona dynamics, we quantified the levels of CENP-E, Spindly and ZW10 throughout early mitosis. This revealed that mid-prometaphase kinetochores in Lis1 depleted cells localised 153±65% more CENP-E when compared to control (Figure 3a). However, this effect was also transient - metaphase kinetochores had equivalent CENP-E levels in siLis1 and siControl treated cells (Figure 3b). In contrast, the kinetochore localisation of Spindly and ZW10 were increased in mid-prometaphase (276±158% and 196±115%, respectively) and metaphase cells depleted of Lis1 (438±178% and 302±185%, respectively) (Figure 3c-f). Importantly, siLis1 treatment had no effect on the kinetochore-association of Mad2, CENP-E, Spindly or ZW10 in nocodazole arrested cells (Supplementary figure 2), suggesting that Lis1 influences their localisation indirectly. One such indirect mechanism is the modulation of corona disassembly kinetics. To investigate this possibility, we imaged prometaphase GFP-Spindly-expressing cells treated with siControl or siLis1 every 30s for 5.5min and quantified the cumulative kinetochore GFP-Spindly intensity at each time point. To enable comparison of siControl and siLis1 conditions, each time point was normalised against the t=0s median of the respective condition. In control cells the cumulative kinetochore GFP-Spindly signal decreased with a t^1/2^ of 120s (Figure 4a,b). The removal of GFP-Spindly from kinetochores slowed in siLis1 treated cells, with a t^1/2^ of 530s (Figure 4, a,b). This resulted in Lis1 depleted kinetochores localising 431±132% more GFP-Spindly at t=330s when compared to control (Figure 4c). Taken together, these data demonstrate that Lis1 is required for timely disassembly of the kinetochore corona.

**Figure 3:**
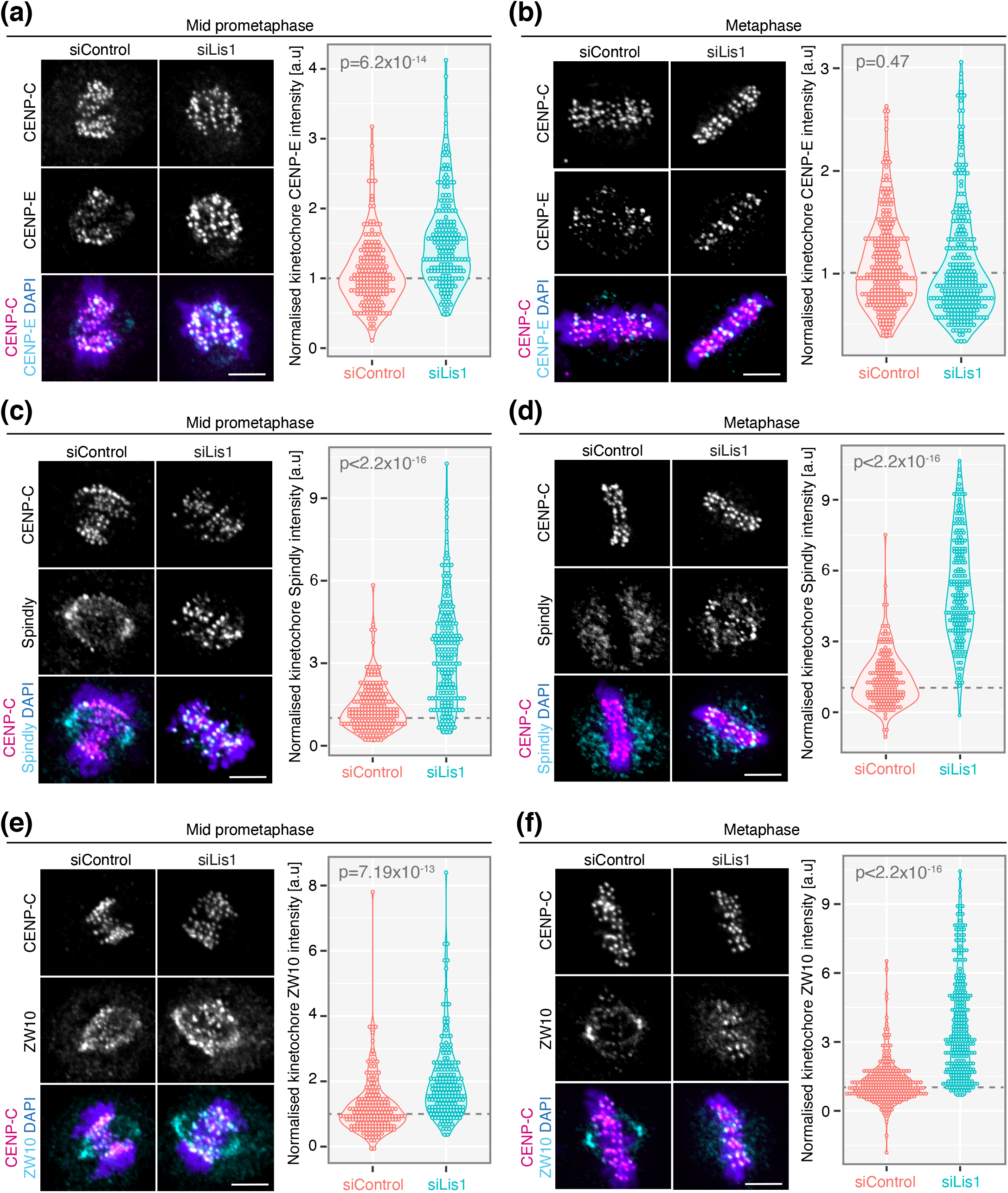
Lis1 is required for complete disassembly of the kinetochore corona. (a) *Left*: Immunofluorescence microscopy images of prometaphase HeLa-K cells treated with siControl or siLis1 and stained with DAPI and antibodies against CENP-E and CENP-C. Scale bar 5*μ*m. *Right*: Quantification of kinetochore CENP-E signal in prometaphase HeLa-K cells treated with siControl or siLis1, n=3, 30 cells, 300 kinetochores per condition. (b) *Left*: Immunofluorescence microscopy images of metaphase HeLa-K cells treated with siControl or siLis1 and stained with DAPI and antibodies against CENP-E and CENP-C. Scale bar 5*μ*m. *Right*: Quantification of kinetochore CENP-E signal in metaphase HeLa-K cells treated with siControl or siLis1, n=3, 30 cells, 300 kinetochores per condition. (c) *Left*: Immunofluorescence microscopy images of prometaphase HeLa-K cells treated with siControl or siLis1 and stained with DAPI and antibodies against Spindly and CENP-C. Scale bar 5*μ*m. *Right*: Quantification of kinetochore Spindly signal in prometaphase HeLa-K cells treated with siControl or siLis1, n=3, 30 cells, 300 kinetochores per condition. (d) *Left*: Immunofluorescence microscopy images of metaphase HeLa-K cells treated with siControl or siLis1 and stained with DAPI and antibodies against Spindly and CENP-C. Scale bar 5*μ*m. *Right*: Quantification of kinetochore Spindly signal in metaphase HeLa-K cells treated with siControl or siLis1, n=3, 30 cells, 300 kinetochores per condition. (e) *Left*: Immunofluorescence microscopy images of prometaphase HeLa-K cells treated with siControl or siLis1 and stained with DAPI and antibodies against ZW10 and CENP-C. Scale bar 5*μ*m. *Right*: Quantification of kinetochore ZW10 signal in prometaphase HeLa-K cells treated with siControl or siLis1, n=3, 30 cells, 300 kinetochores per condition. (f) *Left*: Immunofluorescence microscopy images of metaphase HeLa-K cells treated with siControl or siLis1 and stained with DAPI and antibodies against ZW10 and CENP-C. Scale bar 5*μ*m. *Right*: Quantification of kinetochore ZW10 signal in metaphase HeLa-K cells treated with siControl or siLis1, n=3, 30 cells, 300 kinetochores per condition. Dotted lines indicate the siControl median, to which the data were normalised.

**Figure 4:**
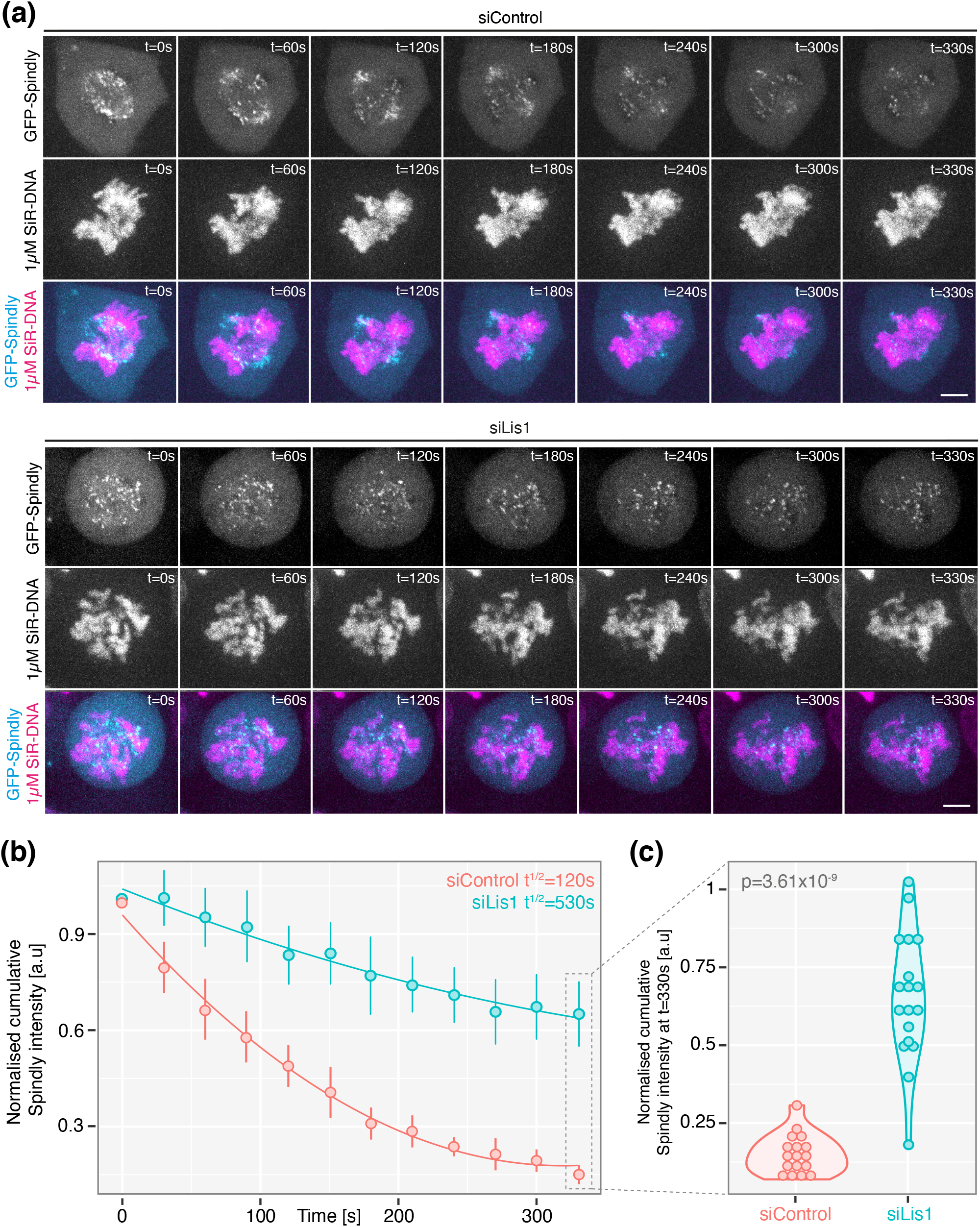
Live-cell analysis of corona dynamics in Lis1-depleted cells. (a) Movie stills of GFP-Spindly expressing HeLa cells treated with siControl or siLis1, imaged every 30s for 5.5min as the cell transited through late prometaphase. Chromosomes were visualised with 1*μ*M SiRDNA. Scale bar 5*μ*m. (b) The cumulative GFP-Spindly intensity for all kinetochores per time point in siControl or siLis1 treated cells imaged every 30s for 5.5min during late prometaphase, n=3, 17 cells siControl, 18 cells siLis1. (c) The cumulative kinetochore GFP-Spindly intensity for all kinetochores in siControl or siLis1 treated cells at the end (t=330s) of the live-cell imaging experiments depicted in (a), n=3, 17 cells siControl, 18 cells siLis1. Points represent individual cells.

### Lis1 and dynein are co-dependent for kinetochore localisation

Thus far, we have established that Lis1 is required for timely and complete disassembly of the kinetochore corona, but via what mechanism? Lis1 depletion could impair the formation and/or maturation of end-on microtubule attachments - the loss of this molecular cue would then indirectly prevent corona compaction. To test this, we quantified the kinetochore proximal α-tubulin signal and kinetochore SKAP signal in siControl or siLis1 treated cells at metaphase. Together, these analyses provide a quantitative readout of K-fibre-kinetochore attachment status (Auckland et al., 2017; Auckland et al., 2020). Crucially, both measures were comparable between siControl and siLis1 treated cells (Supplementary figure 3a-d), suggesting that Lis1 alters corona dynamics via a different mechanism. The minus-end directed motor dynein is essential for the transport of corona cargoes off kinetochores and Lis1 is widely implicated in its localisation. Whether this is conserved at kinetochores is unknown. To address this, we stained siControl or siLis1 treated cells arrested in 3.3*μ*M nocodazole with DAPI and antibodies against dynein intermediate chain (DIC) and CENP-C. Quantification revealed that Lis1 depletion reduced kinetochore DIC localisation by 69±19% when compared to control (Figure 5a,b).

**Figure 5:**
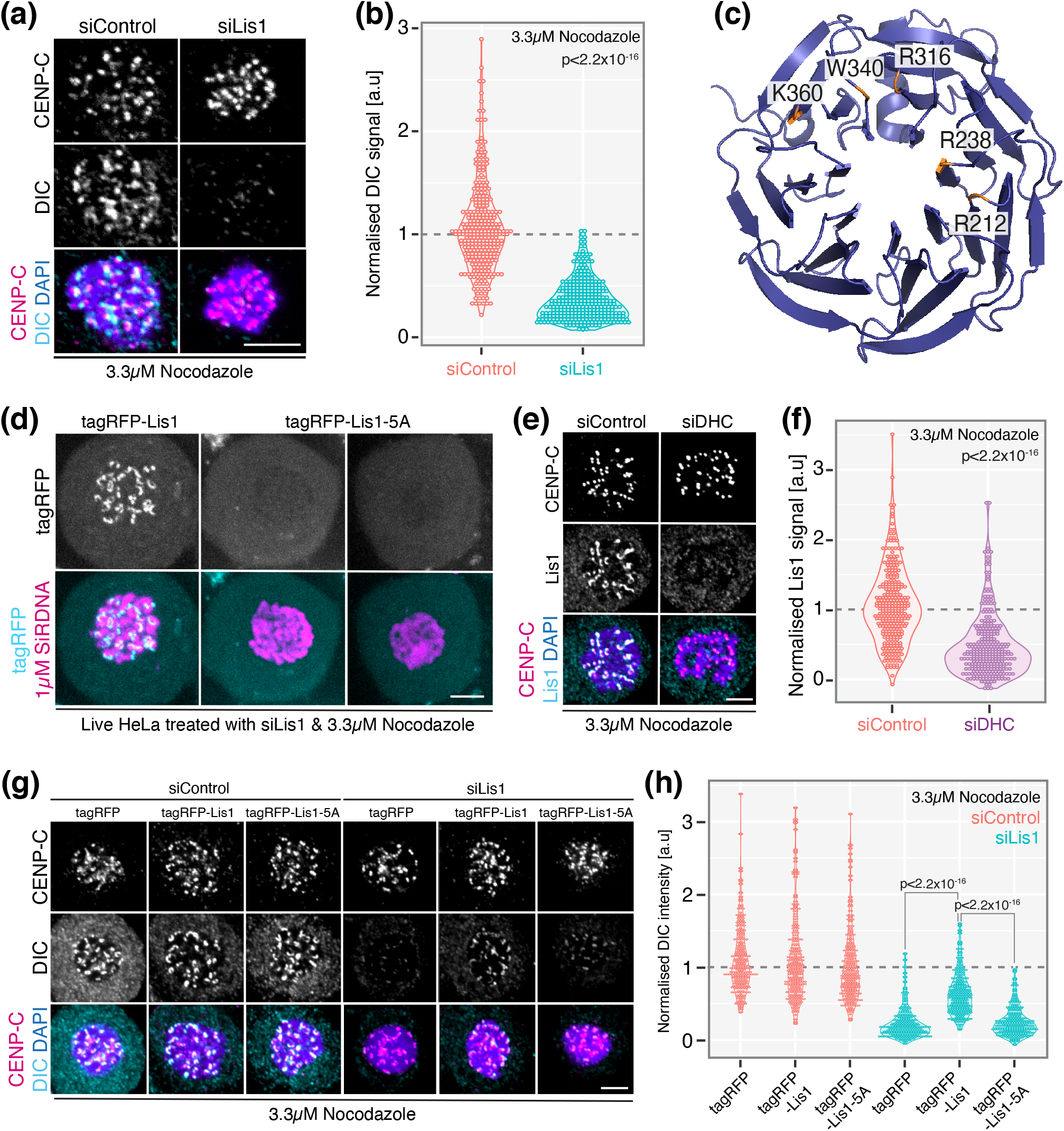
Lis1 and dynein are co-dependent for kinetochore localisation. (a) Immunofluorescence microscopy images of HeLa-K cells treated with siControl or siLis1, arrested in 3.3*μ*M nocodazole for 3hr and stained with DAPI and antibodies against dynein intermediate chain (DIC) and CENP-C. Scale bar 5*μ*m. (b) Quantification of kinetochore DIC signal in HeLa-K cells treated with siControl or siLis1 and arrested in 3.3*μ*M nocodazole, n=3, 30 cells, 300 kinetochores per condition. (c) Ribbon diagram of the Lis1 propeller showing the five residues mutated to alanine in Lis1-5A. (d) Live HeLa-K cells treated with siLis1, transfected with tagRFP-Lis1 or tagRFP-Lis1-5A, and arrested in 3.3*μ*M nocodazole for 3hr. Chromosomes were visualised with 1*μ*M SiRDNA. Scale bar 5*μ*m. (e) Immunofluorescence microscopy images of HeLa-K cells treated with siControl or siDHC, arrested in 3.3*μ*M nocodazole for 3hr and stained with DAPI and antibodies against Lis1 and CENP-C. Scale bar 5*μ*m. (f) Quantification of kinetochore Lis1 signal in HeLa-K cells treated with siControl or siDHC and arrested in 3.3*μ*M nocodazole, n=3, 30 cells, 300 kinetochores per condition. (g) Immunofluorescence microscopy images of the Lis1-5A DIC rescue experiment. Here, HeLa-K cells were treated with siControl or siLis1, transfected with tagRFP, tagRFP-Lis1 or tagRFP-Lis1-5A, arrested in 3.3*μ*M nocodazole and stained with DAPI and antibodies against DIC and CENP-C. Scale bar 5*μ*m. (h) Quantification of kinetochore DIC signal in the Lis1-5A rescue experiment shown in (g), n=3, 30 cells, 300 kinetochores per condition. Dotted lines represent the siControl or siControl+tagRFP median, respectively, to which the data were normalised.

Mutation of five conserved residues on the Lis1 propellor abrogates dynein binding in yeast and severely diminishes binding in humans (Figure 5c, from here on referred to as Lis1-5A) (Htet et al., 2020; Toropova et al., 2014). To further investigate the functional relationship between Lis1 and dynein, we constructed tagRFP-labelled and siRNA-protected Lis1-5A. Firstly, we sought to determine the subcellular localisation of the Lis1-5A mutant. To do so, we transiently expressed tagRFP-Lis1 or tagRFP-Lis1-5A in live cells depleted of Lis1 and arrested in 3.3*μ*M nocodazole (Figure 5d). As expected, tagRFP-Lis1 expanded into crescents that co-localised with SiRDNA (Figure 5d). In contrast, tagRFP-Lis1-5A localisation was entirely cytoplasmic (Figure 5d), demonstrating that a direct interaction between Lis1 and dynein is required to localise Lis1 to kinetochores. In agreement, Lis1 localisation to kinetochores was reduced by 64±43% in cells depleted of dynein heavy chain (DHC) and arrested in 3.3*μ*M nocodazole (Figure 5e,f; Supplementary figure 3e,f). The incomplete loss of Lis1 from kinetochores in DHC depleted cells likely reflects the partial nature of the siRNA (Supplementary figure 3e,f). We then asked if the direct interaction of Lis1 with dynein was required for DIC localisation to kinetochores. Cells were treated with siControl or siLis1, transfected with an empty vector, tagRFP-Lis1 or tagRFP-Lis1-5A, arrested in 3.3*μ*M nocodazole and stained with antibodies against DIC and CENP-C. Western blotting confirmed that our rescue protocol effectively depleted Lis1 and comparably expressed tagRFP-Lis1 and tagRFP-Lis1-5A (Supplementary figure 3g,h). Due to species and molecular weight restrictions, we could not use typical loading controls; nevertheless, a non-specific band at ~50kDa and Ponceau staining (Romero-Calvo et al., 2010) showed comparable protein loading between lanes (Supplementary figure 3g,h). DIC localisation was reduced to 17±17% in siLis1 treated cells transfected with an empty vector (Figure 5g,h). This was rescued to 50±26% by expression of tagRFP-Lis1 (Figure 5g,h). Lis1 depleted cells transfected with tagRFP-Lis1-5A phenocopied cells expressing the empty vector with a kinetochore DIC level of 20±18% (Figure 5g,h), confirming that a direct interaction between Lis1 and dynein is required for dynein localisation to kinetochores. We note that the methanol fixation required for DIC visualisation quenched tagRFP fluorescence, however, the transfection rate was high (determined by live-cell imaging), thus allowing cells to be selected at random. Taken together, these data show that a direct interaction between Lis1 and dynein, mediated by five residues on the Lis1 propeller, is required for the kinetochore localisation of both molecules.

### Cytoplasmic Lis1-dynein can drive corona disassembly

When the Lis1-dynein interaction is abolished, kinetochores fail to load Lis1 and have significantly reduced dynein localisation. To understand the functional significance of this, we quantified the Spindly intensity at metaphase kinetochores in siControl or siLis1 treated cells rescued with an empty vector, tagRFP-Lis1 or tagRFP-Lis1-5A. In agreement with the data presented in Figure 3, kinetochore Spindly staining was increased to 427±297% in Lis1-depleted cells transfected with an empty vector (Figure 6a,b). In contrast, both tagRFP-Lis1 and tagRFP-Lis1-5A restored Spindly removal from kinetochores in Lis1-depleted cells (Figure 6a,b; metaphase kinetochore Spindly intensities of 129±139% and 118±122%, respectively). This raised the possibility that Lis1 contributes to corona disassembly independently of dynein, as the Lis1-5A mutant exhibits reduced dynein-binding activity. To test this, we incubated siControl or siLis1 treated cells with 10*μ*M Dynapyrazole-A (DynpA, a dynein inhibitor (Steinman et al., 2017), see methods for details) for 10min and quantified the Spindly intensity at late prometaphase kinetochores; we reasoned that dynein-driven corona stripping would be maximal at this time as end-on attachments form and mature. If Lis1 had dynein-independent functions at kinetochores, the inhibition of dynein in Lis1-depleted cells would have an additive effect on Spindly localisation. Consistent with DynpA targeting dynein, kinetochore-associated Spindly increased to 195±167% in siControl+DynpA cells when compared to siControl+DMSO (Figure 6c,d). Moreover, we observed that DynpA caused polar accumulation of Spindly in siControl treated cells, a well-established phenotype of dynein dysfunction (Figure 6e) (Howell et al., 2001) In contrast, DynpA treatment had no effect on kinetochore Spindly in Lis1-depleted cells (100±78% siLis1+DMSO and 87±73% siLis1+DynpA; Figure 6c,d). In agreement, polar Spindly was undetectable in Lis1 depleted cells incubated with DynpA (Figure 6e). Together, these data demonstrate that the contribution of Lis1 to dynein localisation and corona disassembly are separable and that cytoplasmic Lis1-dynein can drive corona stripping.

**Figure 6:**
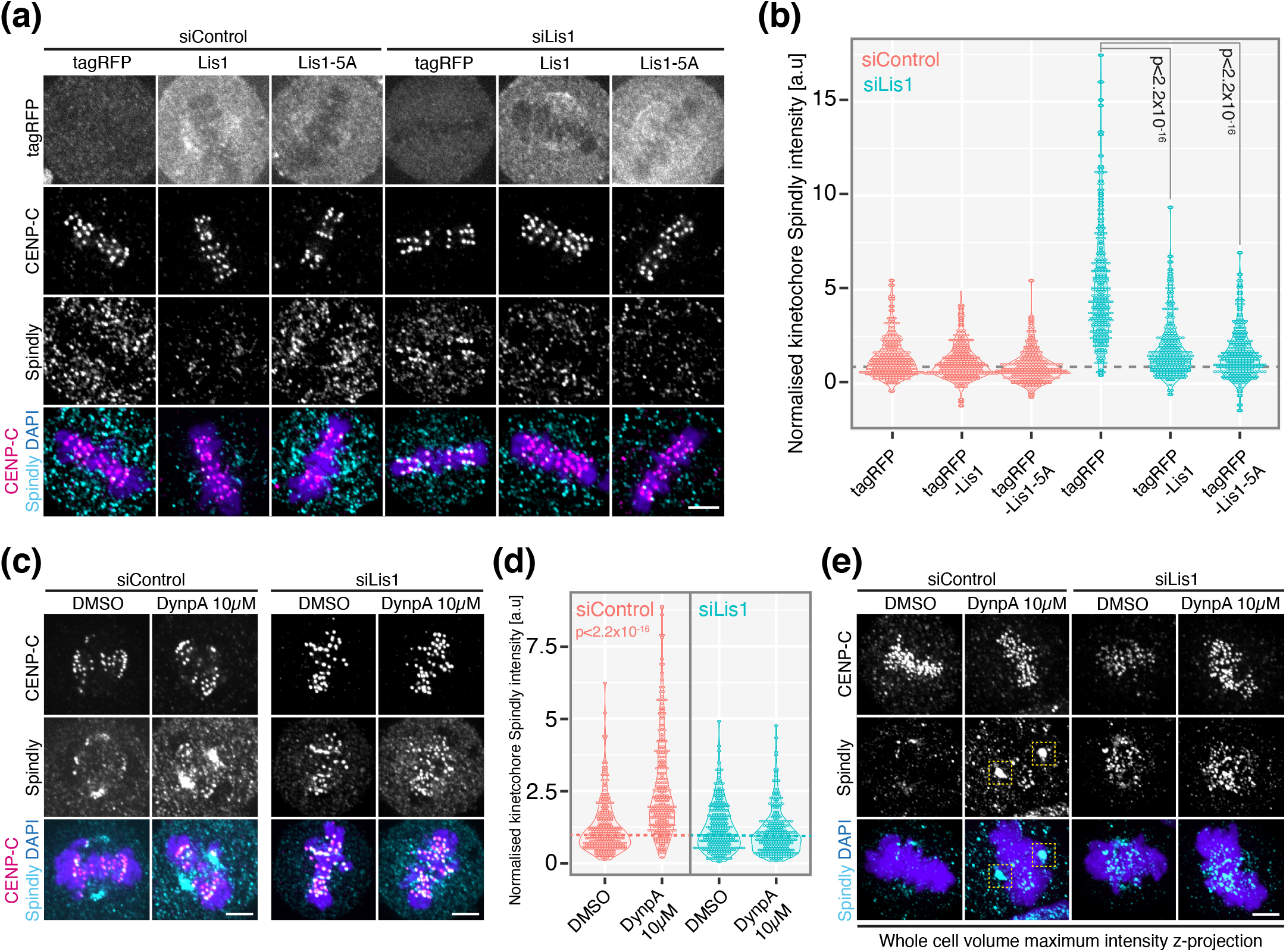
Cytoplasmic Lis1-dynein can drive corona disassembly. (a) Immunofluorescence microscopy images of the Lis1-5A Spindly rescue experiment. Here, HeLa-K cells were treated with siControl or siLis1, transfected with tagRFP, tagRFP-Lis1 or tagRFP-Lis1-5A and stained with DAPI and antibodies against Spindly and CENP-C. Scale bar 5*μ*m. (b) Quantification of kinetochore Spindly signal in the Lis1-5A rescue experiment shown in (a), n=3, ≥29 cells, 290 kinetochores per condition. (c) Immunofluorescence microscopy images of late prometaphase HeLa-K cells treated with siControl or siLis1 incubated with 10*μ*M Dynapyrazole-A for 10min and stained with DAPI and antibodies against Spindly and CENP-C. Scale bar 5*μ*m. (d) Quantification of kinetochore spindly signal in siControl or siLis1 treated cells incubated with 10*μ*M Dynapyrazole-A for 10min, n=3, 30 cells, 300 kinetochores per condition. (e) Whole cell z-projections of siControl or siLis1 treated cells incubated with 10*μ*M Dynapyrazole-A for 10min and stained with DAPI and antibodies against Spindly and CENP-C. Yellow boxes show polar Spindly staining in siControl+Dynapyrazole-A cells. Scale bar 5*μ*m. Dotted lines indicate the siControl+tagRFP, siControl+DMSO or siLis1+DMSO median, respectively, to which the data were normalised.

### Kinetochore-tethered Lis1-dynein is required for error correction

If cytoplasmic Lis1-dynein can drive corona disassembly, why is this module stably tethered to unattached and maturing kinetochores? Dynein forms lateral microtubule attachments that mediate the congression of peripheral chromosomes in early prometaphase (Auckland and McAinsh, 2015; Maiato et al., 2017). However, congression times are comparable between siControl and siLis1 treated cells (Figure 1f,h), suggesting that the loss of Lis1 (and ~69% of kinetochore-associated dynein) does not perturb lateral microtubule capture by kinetochores. In contrast, we found that Lis1-depleted cells exhibited 2.3-fold more anaphase errors than controls (Supplementary figure 4a,b). This is consistent with the finding that dynein prevents the premature stabilisation of erroneous kinetochore-microtubule attachments (Barisic and Maiato, 2015). To explore whether kinetochore-tethered Lis1-dynein is required for attachment error correction, we used a fixed-cell nocodazole release assay. Here, cells were arrested in 330nM nocodazole for 16hr to obtain an almost homogenous population of pseudo-prometaphase cells, thoroughly washed and then incubated for 3.5hr – defects in error correction phenotypically present as cells with misaligned chromosomes. We note that Lis1 depleted cells formed bipolar spindles in monastrol (data not shown) (Raaijmakers et al., 2013), thus preventing the use of the monastrol release assay. In agreement with our live cell imaging, we found that 91±5.1% of siLis1 treated cells expressing an empty vector had misaligned chromosomes 3.5hr after nocodazole release compared with 41±8.2% in siControl+tagRFP (Figure 7a,b). Expression of tagRFP-Lis1 in Lis1 depleted cells rescued chromosome misalignment to 48±4.2% (Figure 7a,b). Crucially, siLis1 treated cells expressing tagRFP-Lis1-5A phenocopied those transfected with an empty vector, with 90±5.2% failing to align all chromosomes (Figure 7a,b). Taken together, these data establish that kinetochore tethered Lis1-dynein is required for the correction of erroneous kinetochore-microtubule attachments.

**Figure 7:**
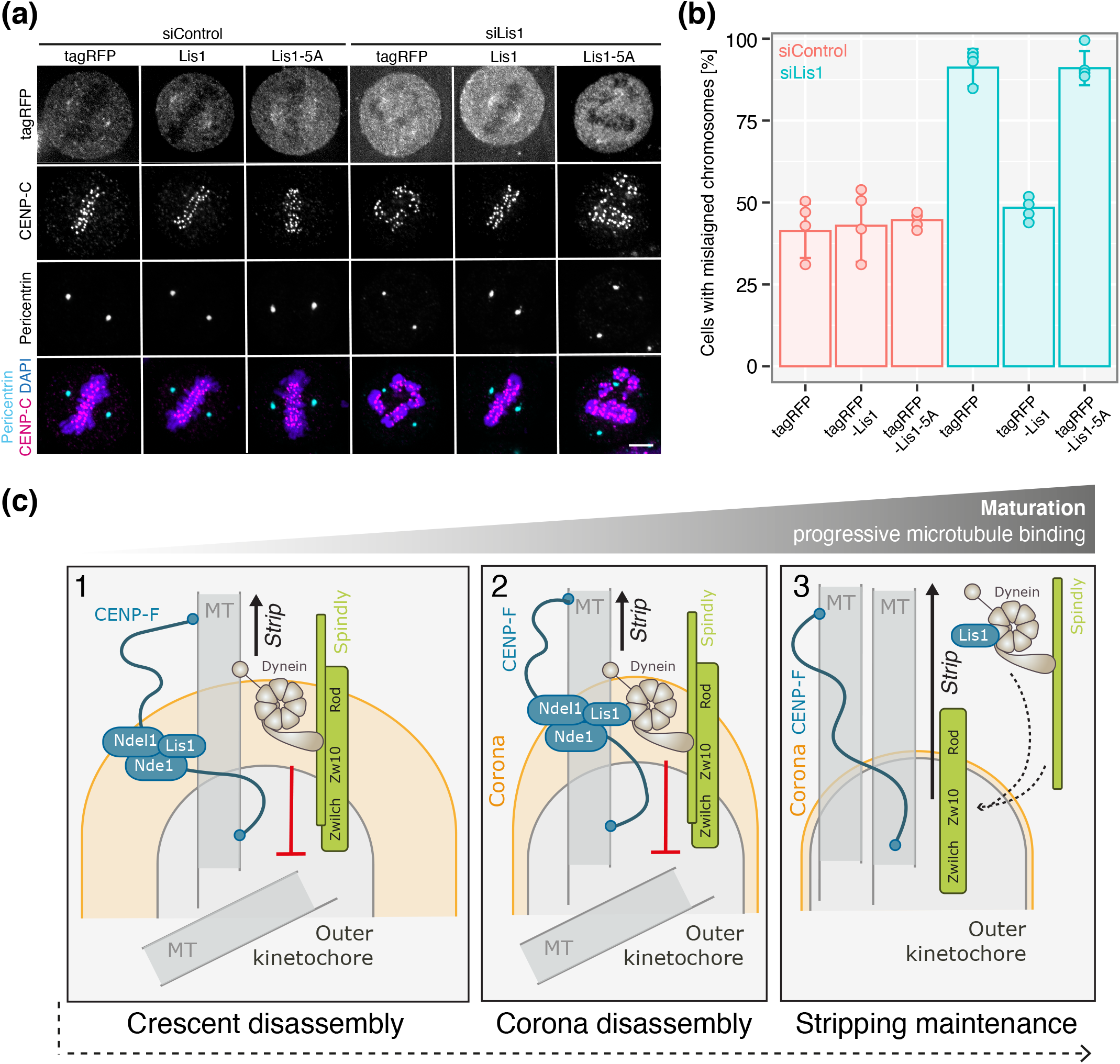
Kinetochore-tethered Lis1-dynein is required for error correction. (a) Immunofluorescence microscopy images of the Lis1-5A error correction rescue experiment. HeLa-K cells were treated with siControl or siLis1 and transfected with tagRFP, tagRFP-Lis1 or tagRFP-Lis1-5A prior to nocodazole washout and staining with DAPI and antibodies against pericentrin and CENP-C. For the nocodazole washout, cells were treated with 330nM nocodazole for 16hr, thoroughly washed with DMEM and incubated for 3.5hr. Scale bar 5*μ*m (b) The proportion of cells in the error correction rescue experiment with misaligned chromosomes 3.5hr after nocodazole release, n=4, ≥88 cells per condition. (c) A model of Lis1 function at the kinetochore. After initial end-on microtubule attachment, Spindly-dynein drives compaction of the maximally expanded crescent-like structure (step 1). Following initial crescent compaction, when end-on attachments are maturing and the corona is still largely associated with kinetochores, Lis1 and Spindly function is coordinated to drive dynein-mediated stripping (step 2). During both phases kinetochore dynein prevents the stabilisation of erroneous attachments (indicated in red). Once attachments have matured at metaphase, where Spindly, dynein and Lis1 are undetectable at kinetochores, cytosolic Lis1-dependent dynein stripping maintains corona compaction (step 3). We suggest that cytosolic Lis1 catalyses the formation of low-level RZZ-Spindly-Dynein-Dynactin complexes, which are templated onto residual kinetochore-associated RZZ, to ensure the removal of corona cargoes as they continually cycle onto metaphase kinetochores.

## Discussion

In this study we have defined for the first time how Lis1 drives corona compaction at kinetochores (summarised in Figure 7c). We suggest that this is a multi-step process where Spindly and Lis1 make multiple contributions dependent upon kinetochore-microtubule attachment maturity. Compaction following initial end-on microtubule binding is driven by Spindly-dynein, as cells expressing a Spindly mutant that abrogates its interaction with dynein fail to disassemble the maximally expanded crescent-like structure (Figure 7c, step 1) (Sacristan et al., 2018). This is distinct from cells depleted of Lis1, where kinetochores form spots that localise excess corona proteins – suggesting that Lis1 acts, at least in part, downstream of Spindly. We note that perturbation of the Spindly-dynein or Lis1-dynein interaction reduce kinetochore dynein by ~75%, respectively (this study and Sacristan et al., 2018), thus it is unlikely that phenotypic differences result from differential effects on dynein localisation. Moreover, microtubule unattached kinetochores formed crescents in the absence of Lis1, therefore defects in crescent formation do not contribute to the phenotypic differences discussed above. Following initial crescent compaction, when end-on attachments are maturing as microtubules bind and the corona is still largely associated with kinetochores, Lis1 and Spindly function is coordinated to drive dynein-mediated stripping (Figure 7c, step 2). This is supported by our observation that Lis1 depletion increased the half-life of kinetochore-associated Spindly in mid-prometaphase. Whether kinetochore associated and/or cytosolic Lis1 is important at this intermediate stage is unclear. Nevertheless, we suggest that following attachment maturation at metaphase, where Spindly, dynein and Lis1 are lost from kinetochores (Supplementary figure 5) (Etemad et al., 2019; Gassmann et al., 2010; Griffis et al., 2007; Monda and Cheeseman, 2018a; Sacristan et al., 2018), cytosolic Lis1-mediated dynein stripping maintains corona compaction. In agreement with this, we find that cytosolic Lis1 can rescue corona disassembly in Lis1-depleted cells at metaphase. How Lis1 and Spindly are integrated here is unclear. However, insight may come from an early study in *Drosophila*, which found that kinetochores replenish low levels of Rod throughout metaphase to maintain dynein-mediated corona stripping (Basto et al., 2004). One possibility is that cytosolic Lis1 catalyses the formation of low-level RZZ-Spindly-Dynein-Dynactin complexes to ensure the removal of corona cargoes as they continually cycle onto end-on attached metaphase kinetochores (Figure 7c, step 3). How these complexes are assembled and how kinetochores transition between initial Lis1-independent crescent disassembly and Lis1 driven compaction maintenance are complex open questions. This is of particular interest given the recent finding that an as yet unknown factor relieves Spindly autoinhibition at kinetochores to promote dynein-dynactin binding (d’Amico et al., 2022).

Our data also provides evidence that corona cargoes are differentially affected by dynein perturbation as cells transition through early mitosis. We find that while the localisation of all tested cargoes is increased at prometaphase kinetochores in Lis1-depleted cells, CENP-E localisation to metaphase kinetochores is unaffected by the loss of Lis1-dynein stripping (unlike Spindly and ZW10). This is consistent with the notion that initial corona compaction is conserved between cargoes and supports the existence of a large complex, such as the RZZ-MES supercomplex (containing Rod, ZW10, Zwlich, Mad1, CENP-E and Spindly) (Samejima et al., 2015), which is stripped by dynein in a single event. However, once kinetochore-microtubule attachments mature, some cargoes are protected from stripping and therefore maintained at metaphase kinetochores. In addition to CENP-E, CENP-F is the other major corona component preserved throughout metaphase (Auckland et al., 2020; Holt et al., 2005; Liao et al., 1995). CENP-E and CENP-F are localised to kinetochores via physical interactions with the dynein-inaccessible receptors BubR1 and Bub1, respectively (Ciossani et al., 2018; Etemad et al., 2019; Gassmann et al., 2010) (see Figure 1a). Thus, as microtubule occupancy increases, the corona transitions into at least two groups with experimentally separable dynamics: (1) Bubs-CENP-E-CENP-F and (2), RZZ-Spindly-Dynein-Mad1-Mad2 (Etemad et al., 2019). We suggest that this allows kinetochores to retain key corona proteins into anaphase while completely shedding SAC components. Indeed, both CENP-E and CENP-F mediate end-on kinetochore-microtubule coupling, an essential requirement for anaphase chromatid segregation (Auckland et al., 2020; Gudimchuk et al., 2013; Vukusic et al., 2019). Future work is needed to fully understand the dynamics of corona disassembly and identify how cargoes transition from ‘strip’ to ‘no-strip’ states in response to microtubule binding. This may depend on post-translational modifications; both CENP-E and CENP-F partially require farnesylation for kinetochore recruitment (Ciossani et al., 2018; Schafer-Hales et al., 2007).

We find that five residues on the Lis1 β-propeller are required for Lis1-dynein localisation to kinetochores and the efficient correction of erroneous kinetochore-microtubule attachments following nocodazole release. Dynein has well-established roles in error correction, where it counteracts the polar ejection force, transports kinetochores towards polar Aurora A, inhibits Rod and modulates Ndc80 microtubule binding. Together, these prevent the stabilisation of erroneous attachments (Amin et al., 2018; Barisic et al., 2014; Cheerambathur et al., 2013; Ye et al., 2015). We reason that the expression of Lis1-5A in Lis1 depleted cells could deregulate these processes by (1) reducing kinetochore dynein localisation by ~75%, (2) altering dynein mechanochemistry such that microtubule and/or Rod binding is perturbed, and/or (3) changing congression dynamics such that kinetochores bypass polar Aurora A. Precisely how mutation of these five residues alter Lis1-dynein binding and/or mechanochemistry is unclear. During preparation of this manuscript the high-resolution structure of the yeast Lis1(Pac1)-dynein complex was reported, which revealed additional dynein contacts in the Lis1 propeller and detailed how the Lis1 stalk interacts with dynein (Gillies et al., 2022). Moreover, this study found that human Lis1 likely contains additional contacts required for dynein regulation. This is consistent with the observation that human Lis1-5A retains some dynein-binding activity whereas yeast Lis1-5A does not {Htet, 2020 #24;Toropova, 2014 #35}. Thus, it is possible that a myriad of complex Lis1-dynein interactions tailor the motor’s functionality. In support of this, the Lis1 stalk is dispensable for microtubule plus-end localisation but necessary for cortical and spindle pole body association in yeast (Gillies et al., 2022). Our data establishes Lis1-5A as another separation-of-function mutant (and the first in humans) that may be a useful tool to further our understanding of Lis1-mediated dynein regulation.

New insights into the pathophysiology of Lissencephaly have come from a study of induced pluripotent stem cell (iPSC)-derived cortical organoids from patients suffering Miller-Dieker syndrome (MDS) (Bershteyn et al., 2017). MDS is characterised by near absent cortical folding (as seen in lissencephalic patients) and is caused by a large heterozygous deletion of human band 17p13.3, which contains dozens of genes including Lis1 (Cardoso et al., 2003; Chong et al., 1997; Dobyns et al., 1991; Dobyns et al., 1983; Reiner et al., 1993). Bersheyn and colleagues found that outer radial glia (ORG) cells had prolonged mitosis and that neuroepithelial stem (NES) cells displayed increased apoptosis (Bershteyn et al., 2017). We suggest our findings provide some mechanistic insight into these phenotypes: (1) prolonged mitosis may result from sustained SAC activation due to perturbed dynein stripping of Mad1-Mad2, and (2) apoptosis may result from catastrophic mitotic errors that program cells for death (Rizzotto et al., 2021). Long time-lapse fate tracking and analysis of kinetochores in ORG and NES cells is needed to build a complete picture of how Lis1 loss-of-function impacts cortical development.

Overall, our experiments have shed light on how Lis1 contextualises dynein function at kinetochores to promote corona disassembly and the correction of erroneous microtubule attachments. Further work combining high-resolution imaging and in vitro reconstitution are needed to fully understand the dynamics of corona disassembly and how corona cargoes engage with dynein. Moreover, it will be essential to determine whether the phenotypes identified here are relevant to other dynein-adapter complexes and cargoes. This is of particular interest given the severe neurodevelopmental defects associated with Lis1 mutations.

## Materials and methods

### Cell culture, siRNA and drug treatments

HeLa-Kyoto (K) and HeLa-LAP-Spindly cells were grown in a humidified incubator at 37°C and 5% CO_2_ in DMEM (Gibco) containing 10% FCS Sigma-Aldrich), 100U/ml penicillin and 100*μ*g/ml puromycin (Invitrogen). siRNA oligonucleotides (53nM, Sigma) were transfected using oligofectamine (Invitrogen) according to the manufacturer’s guidelines and analysed at 48hr (Lis1) or 96hr (DHC). The following sequences were used: control 5’ GGACCUGGAGGUCUGCUGU 3’, Lis1 5’ GAGUUGUGCUGAUGACAAG 3’ and DHC 5’ GGAUCAAACAUGACGGAAU 3’. Drugs were used as follows: nocodazole 3.3*μ*M for 3hr (except for the washout experiment in Figure 7, see below), 100nM or 500nM reversine (see text) and 10*μ*M Dynapyrazole-A for 10min. For Dynapyrazole-A treatment cells were thoroughly washed with warm serum-free DMEM (Gibco) prior to Dynapyrazole-A incubation in serum-free media. For the nocodazole washout, cells were arrested for 16hr in 330nM nocodazole, washed three times with warm DMEM (Gibco) and incubated for 3.5hr before fixation. For overnight imaging, cells were incubated with 1*μ*M SiRDNA (Spirochrome) for 1hr before imaging. For LAP-Spindly induction, cells were incubated with 1*μ*M doxycycline (Sigma-Aldrich) for 24hr before imaging.

### Plasmid construction and siRNA rescue experiments

Full length siRNA-protected Lis1 and Lis1-5A were synthesised (GeneArt), amplified by PCR with flanking SalI and Mfe1 sites and cloned into tagRFP-C1 to create tagRFP-Lis1 and tagRFP-Lis1-5A. The siRNA-protected sequences were as follows: Lis1 5’ GAGCTGCGCAGACGATAAA 3’ and Lis1-5A 5’ GAGCTGCGCAGACGACGCT 3’. For siRNA rescue experiments, cells were seeded on 22mm 1.5 coverglass in DMEM and grown to 50% confluency. Cells were transfected with either siControl or siLis1 using oligofectamine (Invitrogen) in 1.5ml MEM (Gibco) containing 10% FCS (Sigma-Aldrich), 100U/ml penicillin and 100*μ*g/ml puromycin (Invitrogen). After 24hr cells were transfected with 1*μ*g of maxiprep DNA using FugeneHD (Promega) according to the manufacturer’s guidelines in 1.5ml DMEM (Gibco). The media was exchanged for 2mL DMEM (Gibco) 24hr after DNA transfection and the cells incubated for a further 24hr before fixation.

### Immunofluorescence microscopy

Except for DIC and α-tubulin (see below), cells were fixed in 20mM Pipes pH6.8, 10mM EGTA, 1mM MgCl_2_, 0.2% Triton X-100 and 4% formaldehyde for 10min at room temperature (RT). Cells were washed in PBS three times before blocking in PBS containing 3% BSA for 30min and incubation with primary antibodies in blocking solution for 1hr at RT. Primary antibodies are as follows: anti-CENP-C (1/2000, Guinea pig, MBL), anti-CENP-E (1/500, Mouse, Abcam), anti-SKAP (1/400, Rabbit, Atlas Antibodies), anti-Mad2 (1/1000, Mouse, Santa Cruz), anti-Lis1 (1/400, Mouse, Santa Cruz), anti-ZW10 (1/1000, Rabbit, Abcam), anti-Spindly (1/1000, Rabbit, Bethyl), anti-pericentrin (1/1000, Rabbit, Abcam). For anti-DIC (1/200, Mouse, Millipore), cells were fixed in −20°C methanol (Sigma-Aldrich) for 10min and processed as above. The primary antibody was removed with three PBS washes before incubation with Alexa Flour-conjugated secondary antibodies (Invitrogen) at 1/500 for 1hr at RT in blocking solution. Cells were mounted and imaged in Vectashield (Vector Laboratories). For K-fibre analysis, cells were pre-extracted for 30s in 80mM Pipes, pH 6.8, 1mM MgCl_2_, 4mM EGTA, and 0.5% Triton X-100 and fixed by adding glutaraldehyde to 0.5% for 10min. Glutaraldehyde was quenched using a 7min treatment with 0.1% NaBH_4_. Cells were washed three times in PBS for 5min, followed by blocking in TBS containing 0.1% Triton X-100 and 2% BSA for 10min. Cells were then incubated with anti–α-tubulin primary antibody (1/1,000, Mouse, Sigma-Aldrich) for 30min, followed by four 5min washes in TBS containing 0.1% Triton X-100 and incubation with 647nm Alexa Fluor–conjugated secondary antibodies (Invitrogen) in TBS containing 0.1% Triton X-100 and 2% BSA for 30min. Cells were mounted and imaged in Vectashield. 3D image stacks were acquired using a 100X 1.4NA objective on a Yokogawa CSU-W1/Nikon Eclipse Ti2 spinning disk system equipped with an Andor iXon EMCCD camera. Intensities were determined manually in Fiji after background subtraction and normalisation to CENP-C. Conditions were compared using a two-sample t test in R. Distributions were assumed to be normal but this was not formally tested.

### Live-cell imaging

For overnight imaging, cells were seeded in FluroDishes (World Precision) and imaged in DMEM supplemented with 10% FCS (Sigma-Aldrich), 100U/ml penicillin and 100*μ*g/ml puromycin (Invitrogen). 3D image stacks were acquired every 3min for 12hr using a 40X objective on a Yokogawa CSU-W1/Nikon Eclipse Ti2 spinning disk system equipped with an Andor iXon EMCCD camera and Okolab stage top incubator. Mitotic timings and multipolar spindle rates were determined manually in NIS-Elements Viewer (Nikon). Multipolar division rates were compared using a *χ*^2^ test in R. To follow LAP-Spindly dynamics, cells were seeded in FluroDishes (World Precision) and siRNA performed as above. Cells were imaged in DMEM supplemented with 10% FCS (Sigma-Aldrich), 100U/ml penicillin, 100*μ*g/ml puromycin (Invitrogen) and 1*μ*g/ml doxycycline (Sigma-Aldrich). 3D image stacks were acquired every 30s for 5.5min using a 100x 1.4NA objective on the Nikon spinning disk system described above. The sum LAP-Spindly intensities for each time frame were determined manually in Fiji after background subtraction.

### Immunoblotting

Protein extracts were prepared by liquid nitrogen grinding. Briefly, cells were harvested from one 10cm dish and resuspended in 1.5× pellet volumes of H-100 buffer (containing 50mM Hepes, pH 7.9, 1mM EDTA, 100mM KCl, 10% glycerol, 1mM MgCl_2_, and a complete protease inhibitor tablet [Roche]). Cells were ground in liquid N_2_ with a precooled mortar and pestle. The ground cell extract was collected and spun at 14,000 rpm for 30min at 4°C and the soluble fraction collected. Protein concentration was determined by Bradford assay. 15*μ*g of extract was boiled in LDS sample buffer + reducing agent (NuPage) for 10min and separated on a 4–12% Bis-Tris gel (NuPage) in MOPS (NuPage). Proteins were wet-transferred to a nitrocellulose membrane before blocking in 5% milk TBST (Tris-buffered saline with 0.1% Tween 20) for 1hr at RT. The Lis1 primary antibody (1/10,000, Mouse, Santa Cruz) was incubated for 45min at RT in 2% milk TBST. Membranes were washed three times with TBST before incubation with HRP-conjugated secondary antibodies (1/20,000, Abcam) for 45min at RT in 2% milk TBST.

### Figure preparation

All plots were created using ggplot2 in R. Example images were processed in Fiji and figures assembled in Illustrator (Adobe).

## Acknowledgements

We thank A. McAinsh (University of Warwick, Coventry, UK) for reagents, U. Eggert and J. Rosenblatt (King’s College London, London, UK) for sharing equipment and G. Kops (Hubrecht Institute, Utrecht, Netherlands) for the GFP-Spindly cell line. We additionally thank A. McAinsh (University of Wariwck, Coventry, UK) and U. Eggert (King’s College London, London, UK) for critical reading of the manuscript. This work was funded by a King’s Prize Fellowship to PA and a King’s Undergraduate Research Fellowship to OM.

## Author contributions

Project conception, planning, data interpretation and manuscript preparation was carried out by P.A. with contributions from O.M. O.M. conducted the live-cell imaging in Figure 1 and Supplementary figure 4 and constructed the tagRFP-Lis1 and tagRFP-Lis1-5A plasmids. P.W.L. conducted the immunofluorescence experiments shown in Figure 5a,b,e,f and Supplementary figure 3e,f. All other experiments were carried out by P.A.

The authors declare no competing financial interests.

---

**Supplementary figure 1.**
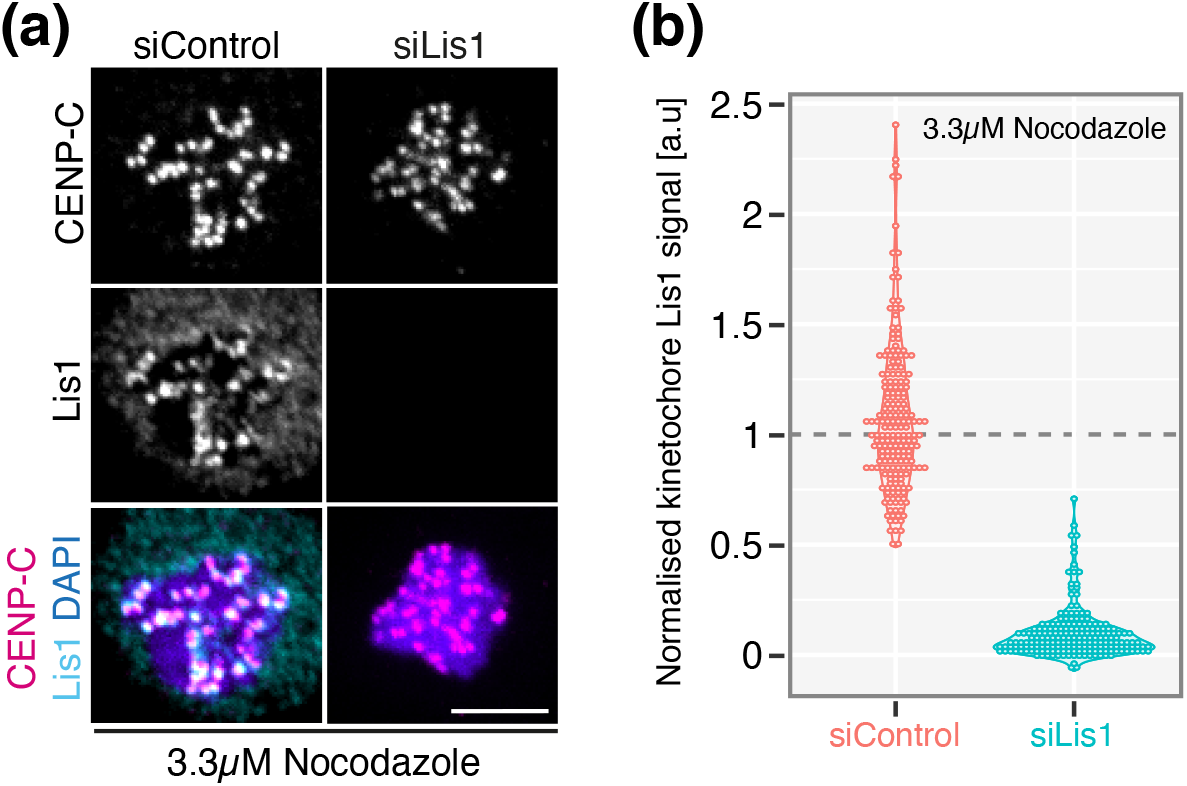
(a) Immunofluorescence microscopy images of HeLa-K cells treated with siControl or siLis1, arrested in 3.3*μ*M nocodazole and stained with DAPI and antibodies against Lis1 and CENP-C. Scale bar 5*μ*m. (b) Quantification of kinetochore Lis1 signal in HeLa-K cells treated with siControl or siLis1 and arrested in 3.3*μ*M nocodazole, n=3, 30 cells, 300 kinetochores per condition. Dotted line represents the siControl median, to which the data were normalised.

**Supplementary figure 2.**
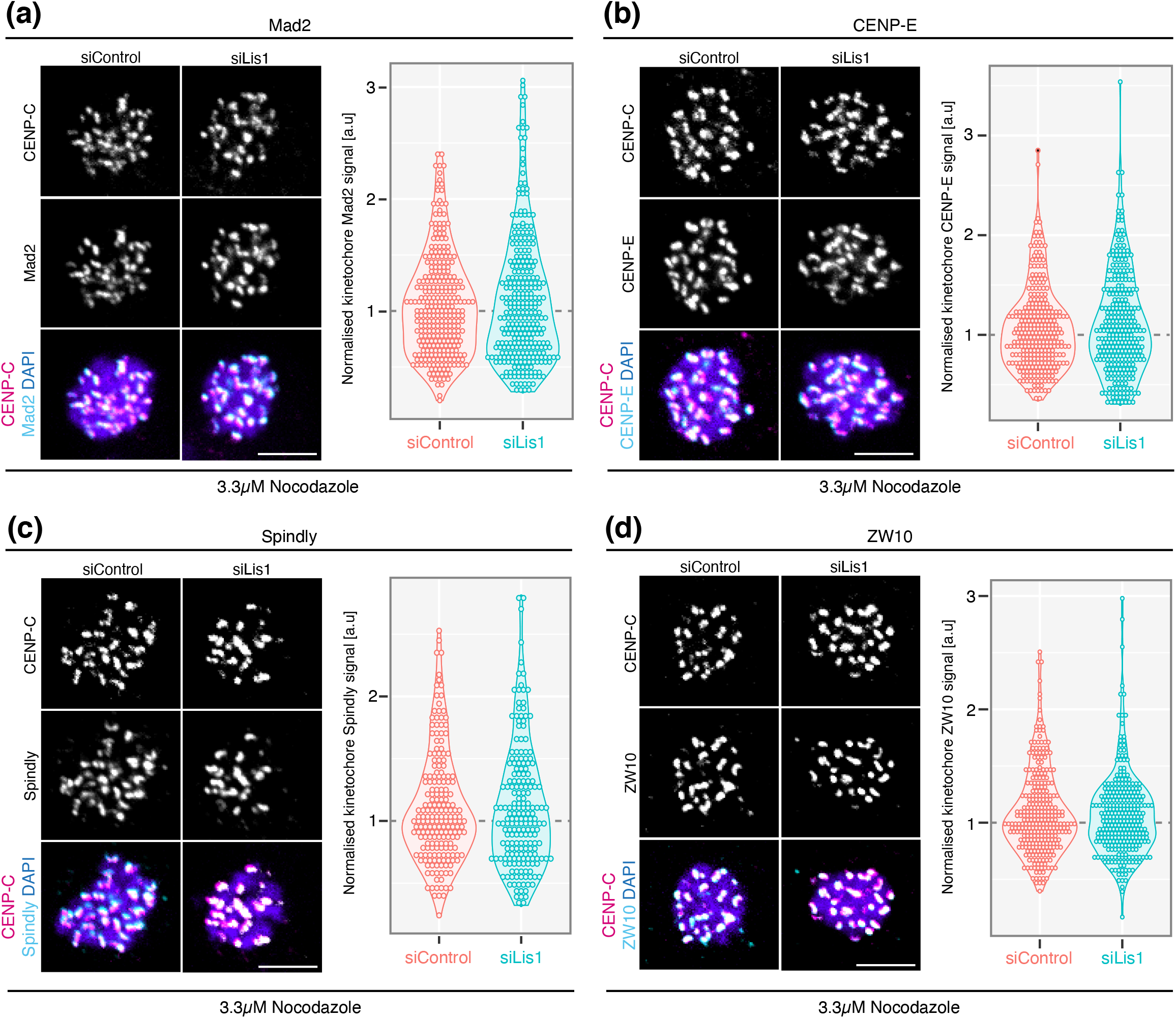
(a) *Left*: Immunofluorescence microscopy images of HeLa-K cells treated with siControl or siLis1, arrested in 3.3*μ*M nocodazole and stained with DAPI and antibodies against Mad2 and CENP-C. Scale bar 5*μ*m. *Right*: Quantification of kinetochore Mad2 signal in HeLa-K cells treated with siControl or siLis1 and arrested in 3.3*μ*M nocodazole, n=3, 30 cells, 300 kinetochores per condition. (b) *Left*: Immunofluorescence microscopy images of HeLa-K cells treated with siControl or siLis1, arrested in 3.3*μ*M nocodazole and stained with DAPI and antibodies against CENP-E and CENP-C. Scale bar 5*μ*m. *Right*: Quantification of kinetochore CENP-E signal in HeLa-K cells treated with siControl or siLis1 and arrested in 3.3*μ*M nocodazole, n=3, 30 cells, 300 kinetochores per condition. (c) *Left*: Immunofluorescence microscopy images of HeLa-K cells treated with siControl or siLis1, arrested in 3.3*μ*M nocodazole and stained with DAPI and antibodies against Spindly and CENP-C. Scale bar 5*μ*m. *Right*: Quantification of kinetochore Spindly signal in HeLa-K cells treated with siControl or siLis1 and arrested in 3.3*μ*M nocodazole, n=3, 30 cells, 300 kinetochores per condition. (d) *Left*: Immunofluorescence microscopy images of HeLa-K cells treated with siControl or siLis1, arrested in 3.3*μ*M nocodazole and stained with DAPI and antibodies against ZW10 and CENP-C. Scale bar 5*μ*m. *Right*: Quantification of kinetochore ZW10 signal in HeLa-K cells treated with siControl or siLis1 and arrested in 3.3*μ*M nocodazole, n=3, 30 cells, 300 kinetochores per condition. Dotted lines represent the siControl median, to which the data were normalised.

**Supplementary figure 3.**
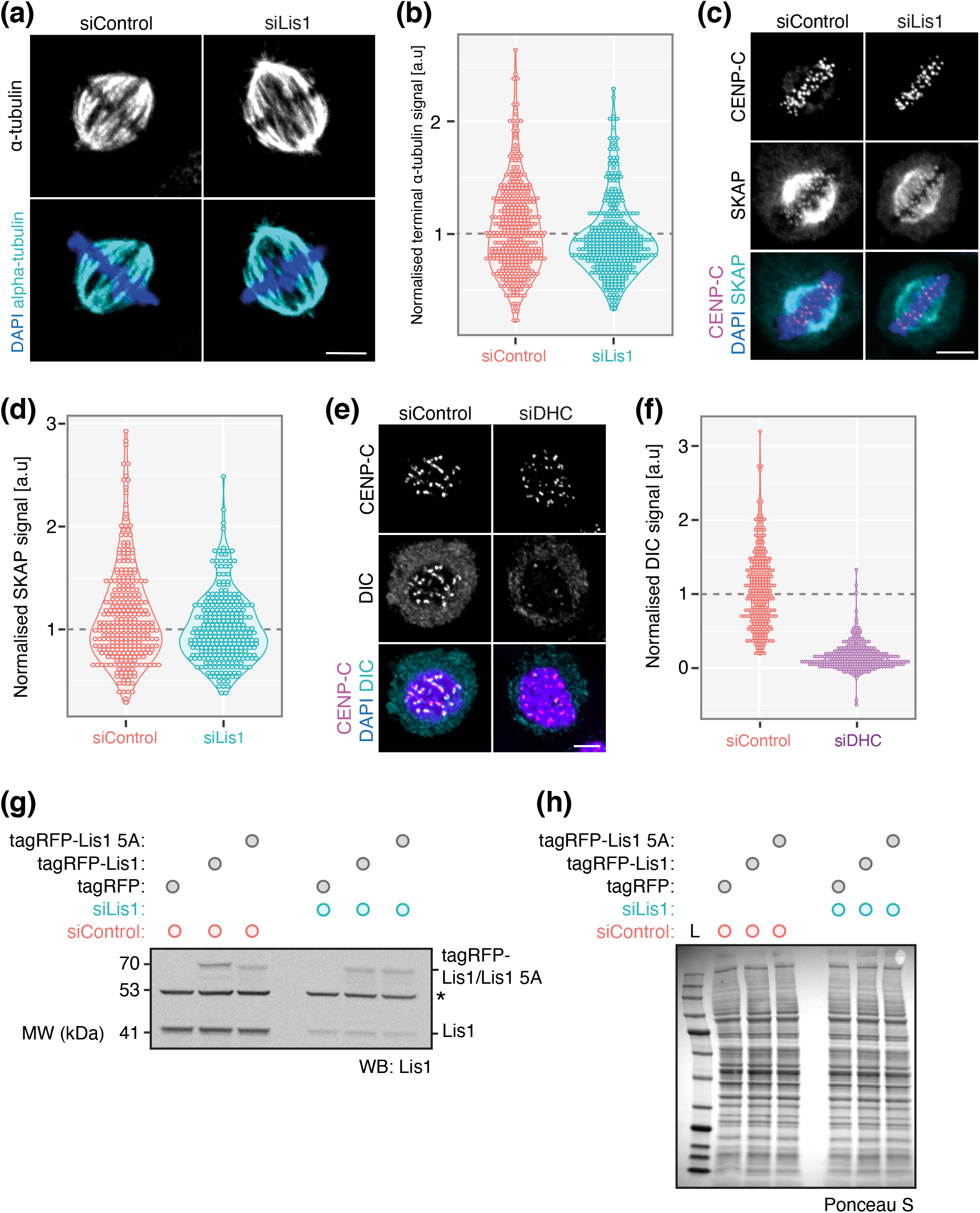
(a) Immunofluorescence microscopy images of HeLa-K cells treated with siControl or siLis1 and stained with DAPI and an antibody against α-tubulin. Scale bar 5*μ*m. (b) Quantification of K-fibre terminal (chromatin proximal) α-tubulin signal in HeLa-K cells treated with siControl or siLis1, n=3, 30 cells, 300 kinetochores per condition. (c) Immunofluorescence microscopy images of HeLa-K cells treated with siControl or siLis1 and stained with DAPI and antibodies against SKAP and CENP-C. Scale bar 5*μ*m. (d) Quantification of kinetochore SKAP signal in HeLa-K cells treated with siControl or siLis1, n=3, 30 cells, 300 kinetochores per condition. (e) Immunofluorescence microscopy images of HeLa-K cells treated with siControl or siDHC, arrested in 3.3*μ*M nocodazole and stained with DAPI and antibodies against DIC and CENP-C. (f) Quantification of kinetochore DIC signal in HeLa-K cells treated with siControl or siLis1 and arrested in 3.3*μ*M nocodazole, n=3, 30 cells, 300 kinetochores per condition. (g) Immunoblot of liquid N_2_ extracts collected from siControl or siLis1 treated cells transfected with tagRFP, tagRFP-Lis1 or tagRFP-Lis1-5A, respectively. The membrane was probed with an antibody against Lis1. Asterisk labels a non-specific band at ~50kDa. (h) Ponseau-S staining of the Lis1 immunoblot membrane in (g). Dotted lines represent the siControl median, to which the data were normalised.

**Supplementary figure 4.**
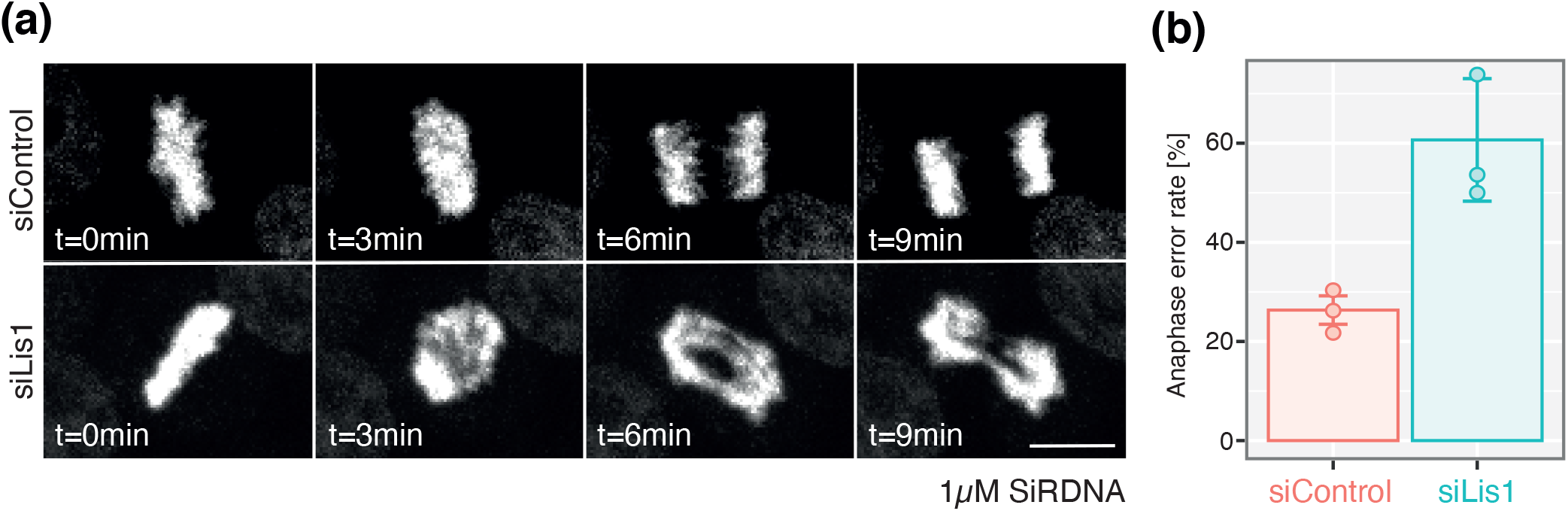
(a) Movie stills of live HeLa-K cells in anaphase treated with siControl or siLis1 and stained with 1*μ*M SiRDNA. (b) Quantification of anaphase error rates in HeLa-K cells treated with siControl or siLis1 and stained with 1*μ*M SiRDNA, n=3, ≥600 cells per condition.

**Supplementary figure 5.**
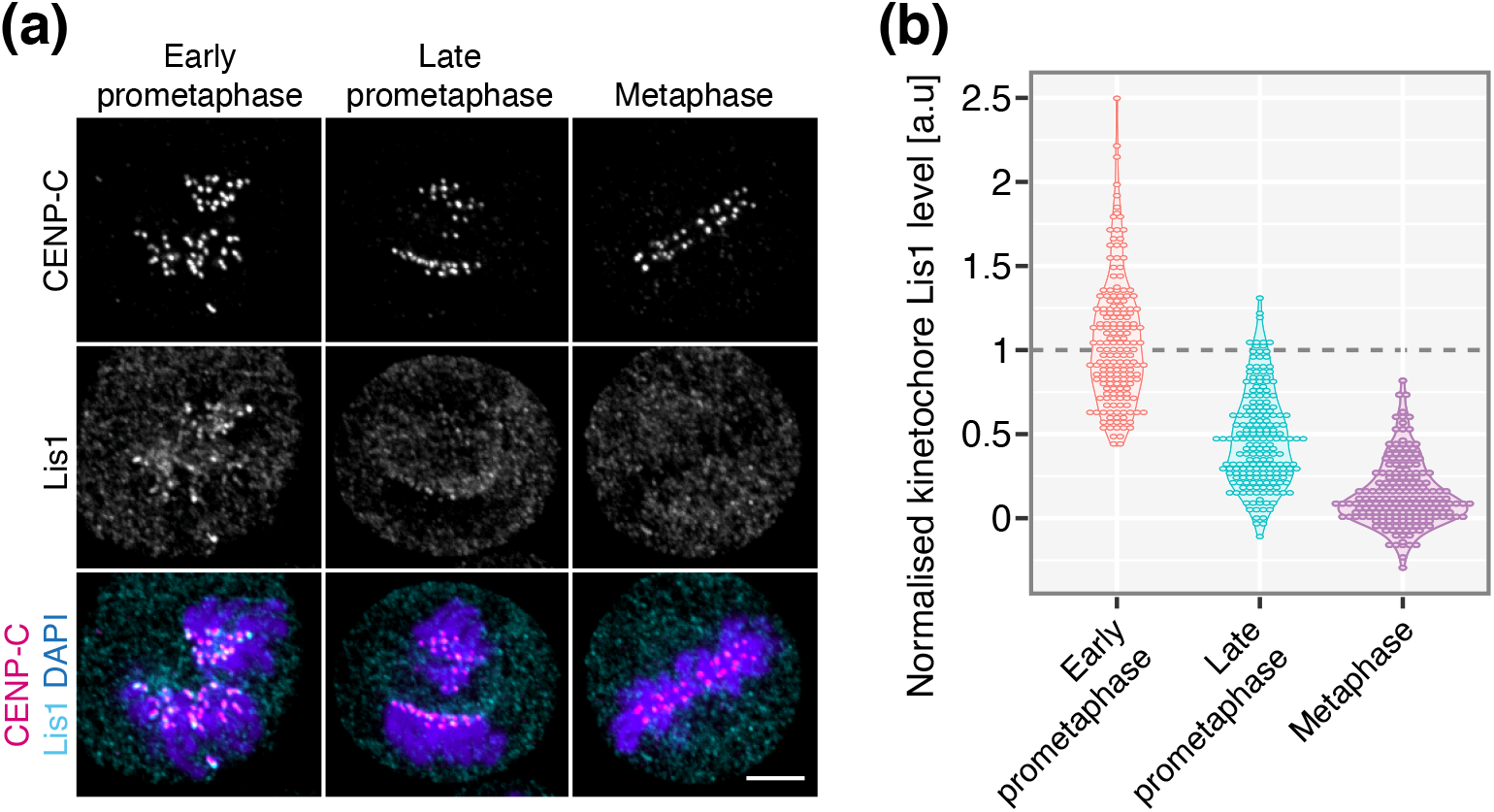
(a) Immunofluorescence microscopy images of early prometaphase, late prometaphase and metaphase HeLa-K cells stained with DAPI and antibodies against Lis1 and CENP-C. Scale bar 5*μ*m. (b) Quantification of kinetochore Lis1 signal in early prometaphase, late prometaphase and metaphase HeLa-K cells. Dotted line represents the early prometaphase median, to which the data were normalised.

